# Characterizing and addressing error modes to improve sequencing accuracy

**DOI:** 10.1101/2024.02.01.578321

**Authors:** Semyon Kruglyak, Andrew Altomare, Mark Ambroso, Vivian Dien, Bryan Lajoie, Kelly N. Wiseman, Shawn Levy, Matthew Kellinger

## Abstract

The accuracy of a sequencing platform has traditionally been measured by the %Q30, or percentage of data exceeding a basecall accuracy of 99.9%. Improvements to accuracy beyond Q30 may be beneficial for certain applications such as the identification of low frequency alleles or the improvement of reference genomes. Here we demonstrate how we achieved over 70% Q50 (99.999% accuracy) data on the AVITI™ sequencer. This level of accuracy required us to not only improve sequencing quality but also to mitigate library preparation errors and analysis artifacts.

## Introduction

Highly parallelized sequencing platforms have revolutionized the ability to generate genome-scale data at costs that support large-scale studies across a diversity of application areas^1–6^. All sequencing platforms share important parameters such as accuracy, read length, and throughput. The combination of parameters defines the capability of the sequencing platform. Read length and throughput are two straightforward parameters to compare between platforms. Accuracy is more challenging to compare due to nuances in the definition and reliance on either predicted quality scores or alignment-based truth sets. This study focuses on accuracy, defined as the concordance between the output of the sequencer and the input DNA. A common measure of accuracy reported by sequencing platform providers is the fraction of data that exceeds an accuracy of 99.9% (commonly referred to as %Q30). Using technology developed by Element Biosciences, we have recently described a new sequencing platform that achieves a high fraction of Q40 data (99.99% accuracy) when applied to PCR-free libraries^7^. Briefly, Element’s avidity base chemistry (ABC) is a cycle-based sequencing method that separates the processes of stepping along the DNA strand that is being sequenced from the process of detecting and resolving each base in the DNA strand. The resolution of each base is achieved through the binding of a labeled polymer termed an avidite. Independent optimization of the stepping and resolving processes with respect to reagent recipes and fit-for-purpose enzymes is an important contributor to ABC accuracy. We demonstrated that accuracy via quality score recalibration of sequencing data, which showed that the quality score assignments above Q40 agreed with empirically determined accuracy^7^. Base quality score recalibration (BQSR) works by considering the assigned base calls and quality scores (along with several covariates) and true calls determined via read alignment to a well-characterized reference genome^8^. If the quality scores assigned by the instrument software are accurate, then they remain unchanged by BQSR. However, if the quality scores overpredict or underpredict the accuracy of the underlying data, then they are modified accordingly in the BQSR step. We decided to use BQSR as a means of evaluating additional optimizations to the sequencing recipe. As sequencing accuracy improved, we expected to observe increases in recalibrated Q scores. In our attempt to further improve the accuracy of ABC sequencing, the percentage of bases exceeding Q30 and Q40 increased but the highest qualities achieved via base quality recalibration did not substantially improve. The results appeared to show that ABC hit a sort of Q-score ceiling.

Here we describe the cause of this apparent Q ceiling and recent advances to the Element platform and ABC sequencing that have led to 70% of sequencing data exceeding Q50 or 99.999% accuracy. Demonstrating this level of accuracy involved reevaluating each step of the DNA sequencing process, including library preparation, ABC sequencing, and data analysis. A key finding was that the majority of high-quality errors (i.e. bases that were assigned a high quality score but mismatched the reference) come from processes either upstream or downstream of sequencing. We first characterized and quantified errors associated with each step. We then implemented improvements that reduced the number of errors from the primary contributors. We leveraged model systems to enhance our understanding of various error processes and to develop improvements. We then applied the optimized protocols to human whole genome sequencing and observed significantly reduced mismatch rates.

## Methods

To improve sequencing accuracy, we optimized the ABC sequencing recipe. Optimizations included modifying reagent concentration, contact times, and new enzymes. The results showed improvements to primary sequencing metrics such as phasing and signal to noise, as well as to average mismatch rate. We did not observe significant increases in the highest recalibrated quality scores. We therefore hypothesized that processes upstream or downstream of sequencing were affecting the recalibration. For example, if an error occurs during library preparation (e.g. damage during DNA fragmentation causing a C to T mismatch) then the sequencer is likely to call a high-quality T at the position because an error occurring in library preparation would not affect any predictors of sequencing quality such as intensity or phasing rate. However, following alignment, the base would not match the reference, so the recalibration process would associate the error with the high-quality bin. If the frequency of such errors exceeds 1 in 100,000, then the quality bin will recalibrate below Q50. In addition to library preparation, analysis artifacts can also generate high-quality errors that decrease recalibrated quality scores. Examples include incorrect alignments causing differences between the published reference genome and the DNA being sequenced. The frequency of analysis errors is expected to increase with the complexity of the reference genome. To characterize analysis errors, we sequenced the PhiX and *E. coli* genomes at high coverage. These model systems facilitate error tabulation because (1) the genomes are haploid, so heterozygous variants are not a concern, (2) the reference genomes are well established allowing accurate error determination compared to a truth set, and (3) alignment of short reads to these genomes is simplified by relatively few repeat regions.

To distinguish library preparation errors and analysis errors from sequencing errors, we focused on short insert lengths, where read 1 (R1) and read 2 (R2) fully overlap. Following alignment and in the absence of any errors, we expect R1 to agree with R2 and both to agree with the reference at every position. (Note that although R1 and R2 are sequenced in opposite directions, both are presented in the forward orientation following alignment.) When considering base calls with high quality scores, disagreements between R1 and R2 are almost certainly sequencing errors because the same molecule is being sequenced twice. Conversely, disagreements with the reference, when high quality calls in R1 and R2 agree with each other are most likely the results of library preparation errors or analysis artifacts, given that the probability of two high-quality sequencing errors at the same position is extremely low. For each 2×150 base pair sequencing run, we aligned the reads with BWA-mem version 0.7.8 with default settings. We then selected reads with insert lengths between 90 and 150 base pairs to ensure complete overlap between R1 and R2. We considered only primary alignments and ignored reads with indels. Indels complicate the comparison of overlapping reads and given that our indel rate is in the order of 10^−6^ to 10^−7^, their contribution is small compared to mismatch errors. We then used a custom python script to compare R1, R2, and the reference sequence to tabulate the different types of errors.

To distinguish library preparation errors from analysis artifacts, we identified regions in the *E. coli* genome where alignments were unreliable or where the reference genome did not accurately represent the input DNA that we sequenced. Specifically, we used GenMap with k-mer size 30 to identify regions of low mappability^9^. We then identified 10 additional sites in the *E. coli* genome where multiple forward and reverse reads in multiple sequencing runs in both Element sequencing and an orthogonal sequencing technology agreed with each other but disagreed with the reference. In total we identified 2.5% of the *E. coli* reference genome expected to be highly enriched for analysis artifacts but not other types of errors. Any read overlapping with the problematic reference regions was removed from error analysis, retaining over 95% of the read data. To assess the impact of the change, we compared the *E*. coli results with and without the modified reference genome, to results from PhiX, which is expected to have the same rate of library preparation errors but virtually no analysis artifacts.

We iterated modifications to library preparation and sequencing recipe with error type characterization to maximize recalibrated quality scores. The recipe combining all the changes is termed UltraQ. Once the sequencing recipe is finalized, we will train new quality tables and produce quality scores based on a set of predictors that are expected to match the recalibrated values.

## Results

Table 1 compares the current version of Element’s ABC sequencing chemistry (termed Cloudbreak) to an optimized recipe, in terms of phasing rate, %Q30, %Q40, and average reference mismatch rate. Although the metrics are significantly improved, the benefits observed in the optimized recipe did not translate to a large increase in the maximum recalibrated quality score.

**Table 1:** Comparison of sequencing metrics between current chemistry and optimized recipe. The average mismatch rate is dominated by a small fraction of low-quality base calls.

| Metrics derived from *E. coli* sequencing | Current chemistry version | Optimized chemistry |
| --- | --- | --- |
| Per cycle phasing | 0.15% | 0.08% |
| Per cycle prephasing | 0.04% | 0.01% |
| Recalibrated %Q30 | 96.7% | 98.5% |
| Recalibrated %Q40 | 88.2% | 96.3% |
| Average mismatch | 0.128% | 0.056% |

We therefore began to characterize the different error types as described in the methods section. To quantify the analysis errors, we sequenced pooled libraries of PhiX and *E. coli* on the same flowcell with the optimized sequencing recipe. The rationale was that the PhiX reference is very well established and alignment to the short PhiX genome is straightforward. PhiX should therefore have almost no errors that are caused by analysis artifacts. Table 2 compares error rates of PhiX and *E. coli*, where R1 and R2 agree with each other, both have high quality, but disagree with the reference. The PhiX rate of such errors corresponds to Q48.4, whereas the *E. coli* rate corresponds to Q45.5, which is roughly a 2-fold difference. To determine whether this difference could be explained by analysis errors, we removed roughly 2.5% of the *E. coli* genome that included low mappability regions and sites where the published reference did not correctly reflect the input DNA (see methods for how these regions and sites were determined). Notably, the sites we removed did not indicate being more difficult to sequence in that they were not enriched for errors where R1 and R2 disagreed with each other. Following reanalysis with the modified reference, the frequency of such errors in *E. coli* corresponded to Q47.6, more closely matching PhiX.

**Table 2:** Comparison of library preparation and analysis errors for a combined E. coli and PhiX sequencing run. The modification to the E. coli reference brings the errors in line with PhiX, suggesting that most of the analysis-specific artifacts are eliminated with the modification. The base comparisons are based on a representative subsample of the run data.

| Metric | PhiX | *E. coli* | *E. coli* – modified reference |
| --- | --- | --- | --- |
| Base comparisons | 75,760,190 | 63,588,653 | 60,822,045 |
| R1 and R2 agree, but differ from reference | 1,096 | 1,776 | 1,057 |
| Implied Q score | Q48.4 | Q45.5 | Q47.6 |

Figure 1 shows examples of analysis artifacts removed by the reference modifications. For subsequent analyses of *E. coli*, we used the modified reference. The contribution of analysis errors is expected to be much higher in the human genome, but efforts like the T2T reference and the Q100 HG002 aim to address such errors. In the meanwhile, we focus on *E. coli*, acknowledging that it does not have the sequence context complexity or the same difficulty present in the human genome.

**Figure 1:**
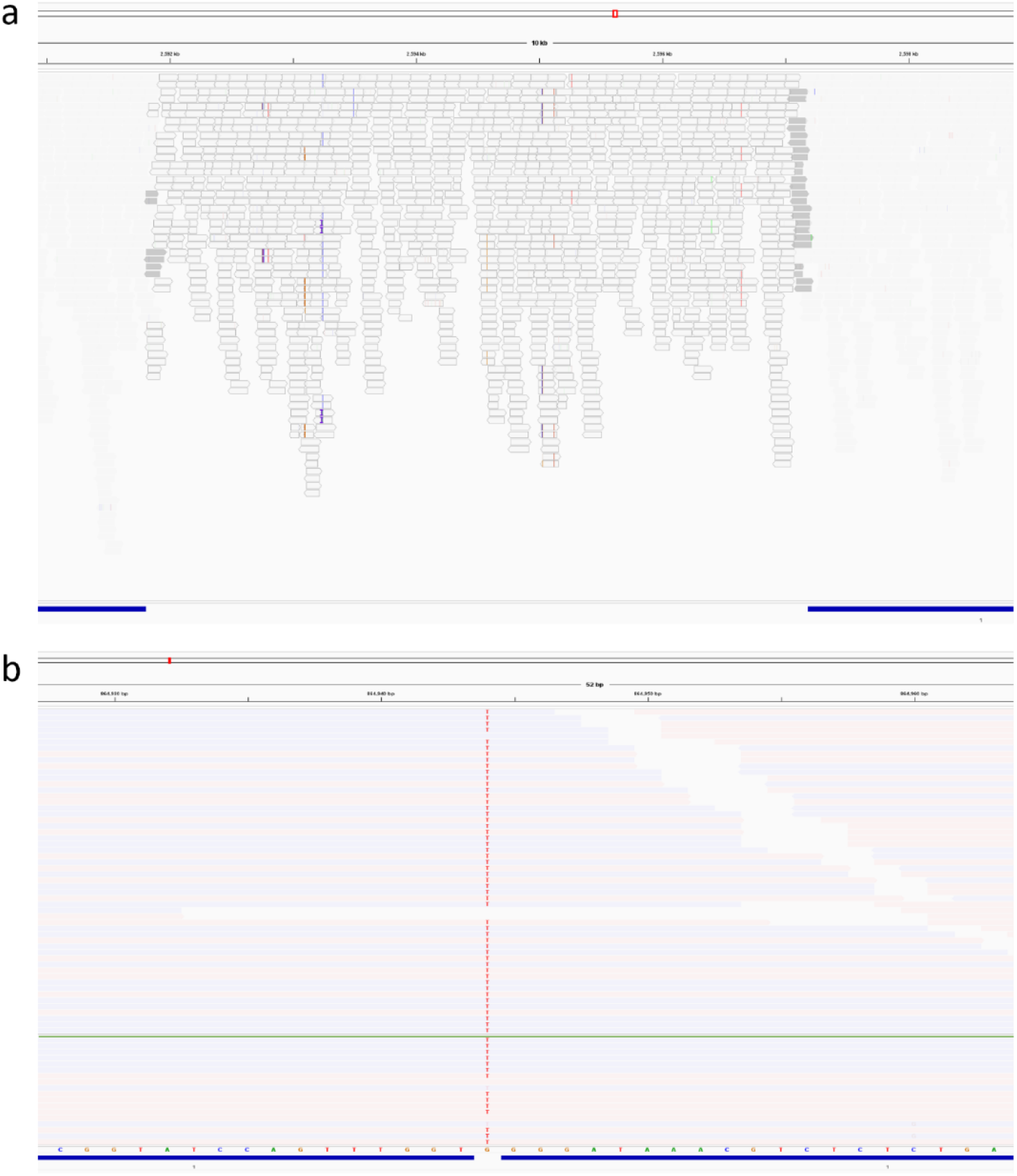
IGV (Integrative Genomic Viewer) displays of errors created by analysis artifacts. (a) shaded reads reflect low mapping quality and colored vertical lines show mismatches resulting from poor alignment. The bar at the bottom indicates portions of the reference retained (blue) and portions excluded (white) as described in methods. (b) Single position that is G according to the reference, but T based on high quality calls from both strands (pink and blue). Reads below the green line were generated by NextSeq sequencing of the same library. The evidence suggests that the published reference is not correct for the sequenced strain.

Our next objective was to distinguish and quantify sequencing errors and library preparation errors by studying short insert lengths as described in the methods section. The first column of table 3 shows the quality scores based on the number of all mismatches, the number of high quality sequencing errors, and the number of library preparation errors from sequencing a different *E. coli* library. Sequencing errors dominate (Q32.7) but they are nearly eliminated when only bases with quality score Q40 and above are considered (empirical Q57 when conditioning on predicted Q>39). By contrast, library preparation errors, which cannot be stratified by quality score, corresponded to Q46 (i.e. more than 10 times the high Q sequencing errors). Although library preparation errors are a small fraction of overall sequencing error, when considering only high-quality calls, they are dominant and limit the recalibration.

**Table 3:**
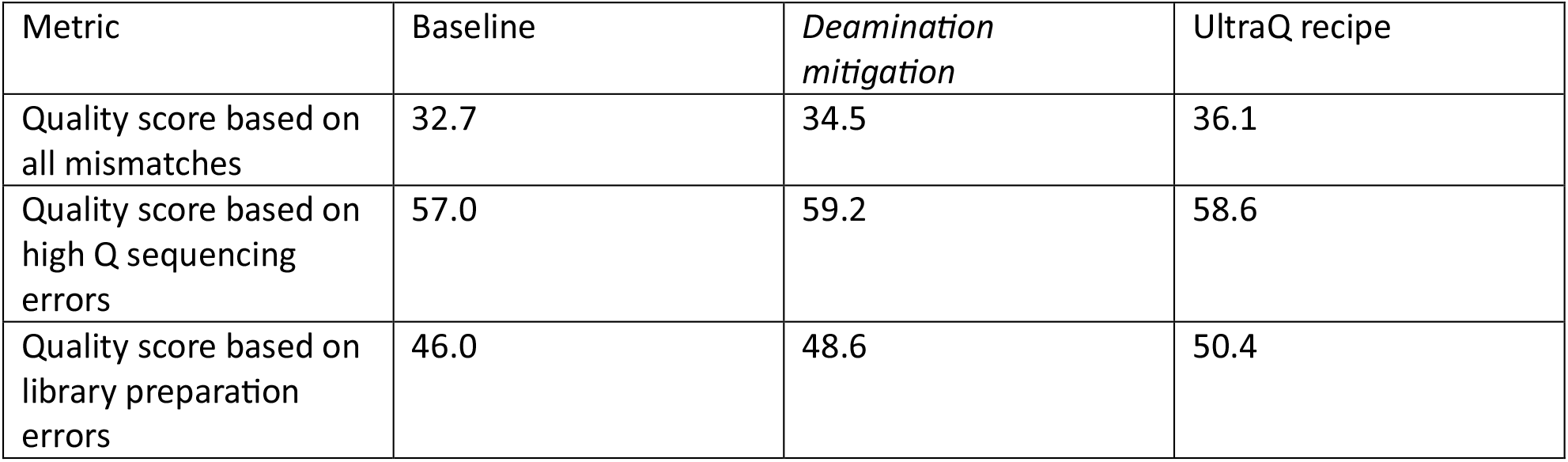
Breakdown of errors by category based on a subset read pairs from each of 3 sequencing runs. Above Q39, the number of sequencing errors is insignificant relative to the number of library prep errors. The UltraQ recipe increases the percent Q40 and dramatically reduces the number of library preparation errors.

| Metric | Baseline | *Deamination mitigation* | UltraQ recipe |
| --- | --- | --- | --- |
| Quality score based on all mismatches | 32.7 | 34.5 | 36.1 |
| Quality score based on high Q sequencing errors | 57.0 | 59.2 | 58.6 |
| Quality score based on library preparation errors | 46.0 | 48.6 | 50.4 |

Having identified library preparation errors as an important contributor, we next attempted to understand and reduce these errors. One well known source of library preparation error is base deamination^10^. This type of DNA damage modifies cytosine to uracil and results in a C to T mismatch in DNA sequencing. We developed a protocol to minimize errors stemming from deamination damage. The protocol leveraged milder conditions for DNA denaturation and introduced a USER enzyme (NEB) treatment that cut library fragments containing a uracil base. Figure 2 shows a comparison of library preparation errors in a baseline run and a run where the deamination protocol was utilized. The errors are normalized by the total number of base pairs compared to the reference. Although the run utilizing the deamination protocol is better overall, the primary difference is the sharp drop in C to T and G to A mismatches. Each of these mismatch signatures can be caused by deamination damage, depending on which DNA strand is sequenced. The second column of table 3 shows the impact on library preparation errors in terms of Q score.

**Figure 2:**
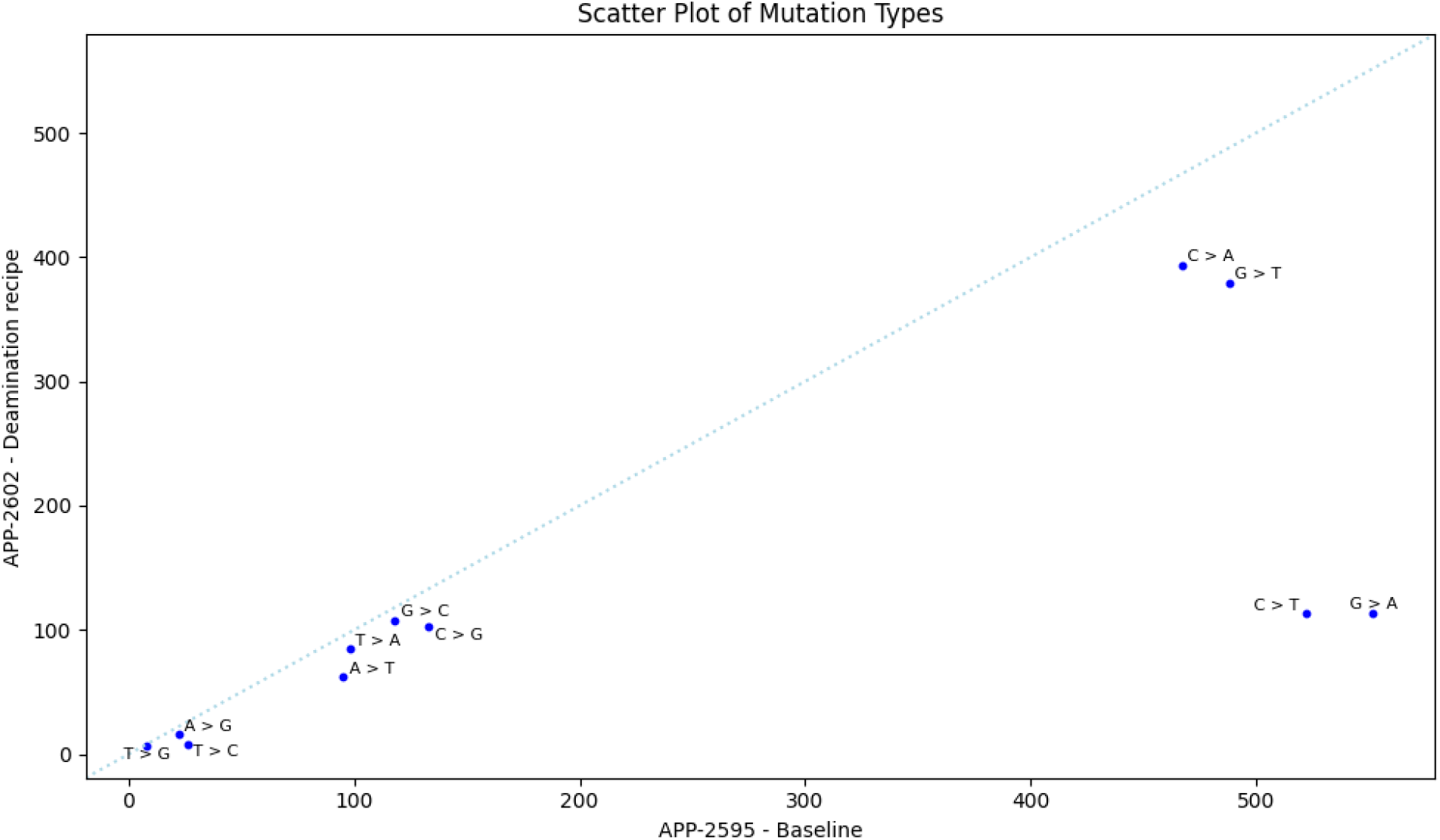
Normalized mismatch counts for two runs. APP-2595 utilizes standard library preparation, while APP-2602 leverages a protocol to minimize reads with deamination damage. APP-2602 is slightly higher quality across mismatch types but shows a large decrease in C > T mismatches (or G > T on the opposite strand).

Another frequent library preparation error signature is G to T (or C to A on the reverse strand). We observed that most such errors occurred in the early cycles of R2. We hypothesize that the errors occur during the end-repair step of library preparation. Due to the 5’ to 3’ direction of DNA replication, blunt ends are created either via exonuclease activity or via gap filling extension. The process of ligating different primers on the two ends ensures that R2 will always start at the end where gap-fill was used. The high frequency of G to T library errors in the fragment sequenced early in R2 suggests that the gap fill process is relatively low fidelity. While we consider methods to address this issue, one way to sidestep the gap fill error is to skip the early bases of the fragment where they frequently occur. We performed dark cycling^†^ during the initial 15 bases in the second sequencing read. Dark cycling effectively skips the problematic portions of the fragment, with only a small increase to in the total run time. The run also leveraged the deamination protocol in library preparation. Figure 3 shows the impact on all library preparation error types. The G to T and C to A errors are sharply decreased and all library preparation errors are relatively infrequent. In this run, the library preparation error correction resulted in Q50.4 accuracy. The sequencing errors are even less frequent (Q58.6) but library preparation errors no longer prevent us from achieving Q50 (see third column of table 3).

**Figure 3:**
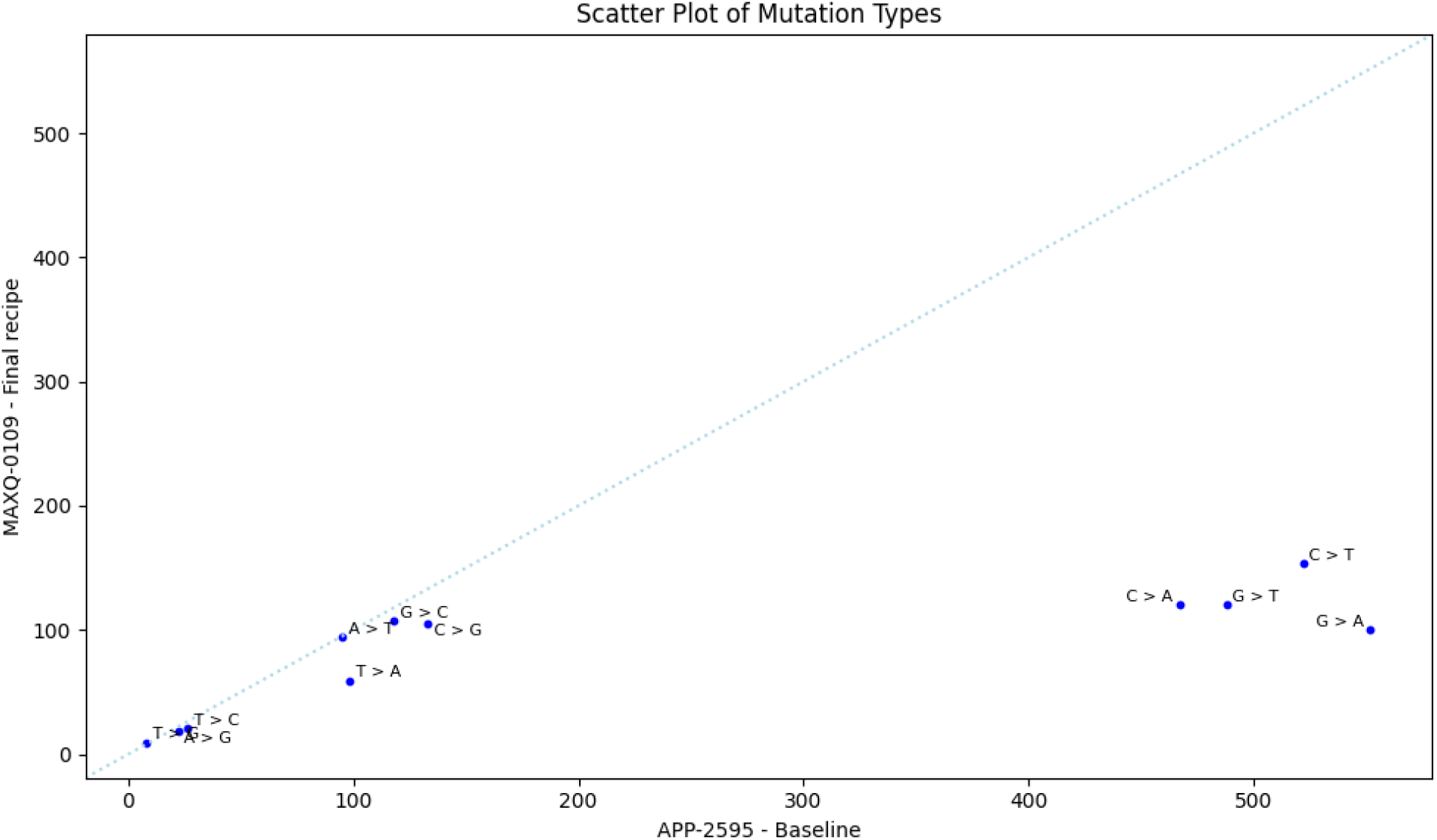
Normalized mismatch counts for a standard sequencing (APP-2595) and a run that uses both the recipe to minimize deamination and dark cycling through the first 15 bp of R2. Both the C>T (G>A) and G>T (C>A) error types are significantly reduced and in line with other error types.

Figure 4 shows a quality score recalibration for the entire run, without the short insert length constraint. The percent recalibrated Q50 for the run is 78.1%.

**Figure 4:**
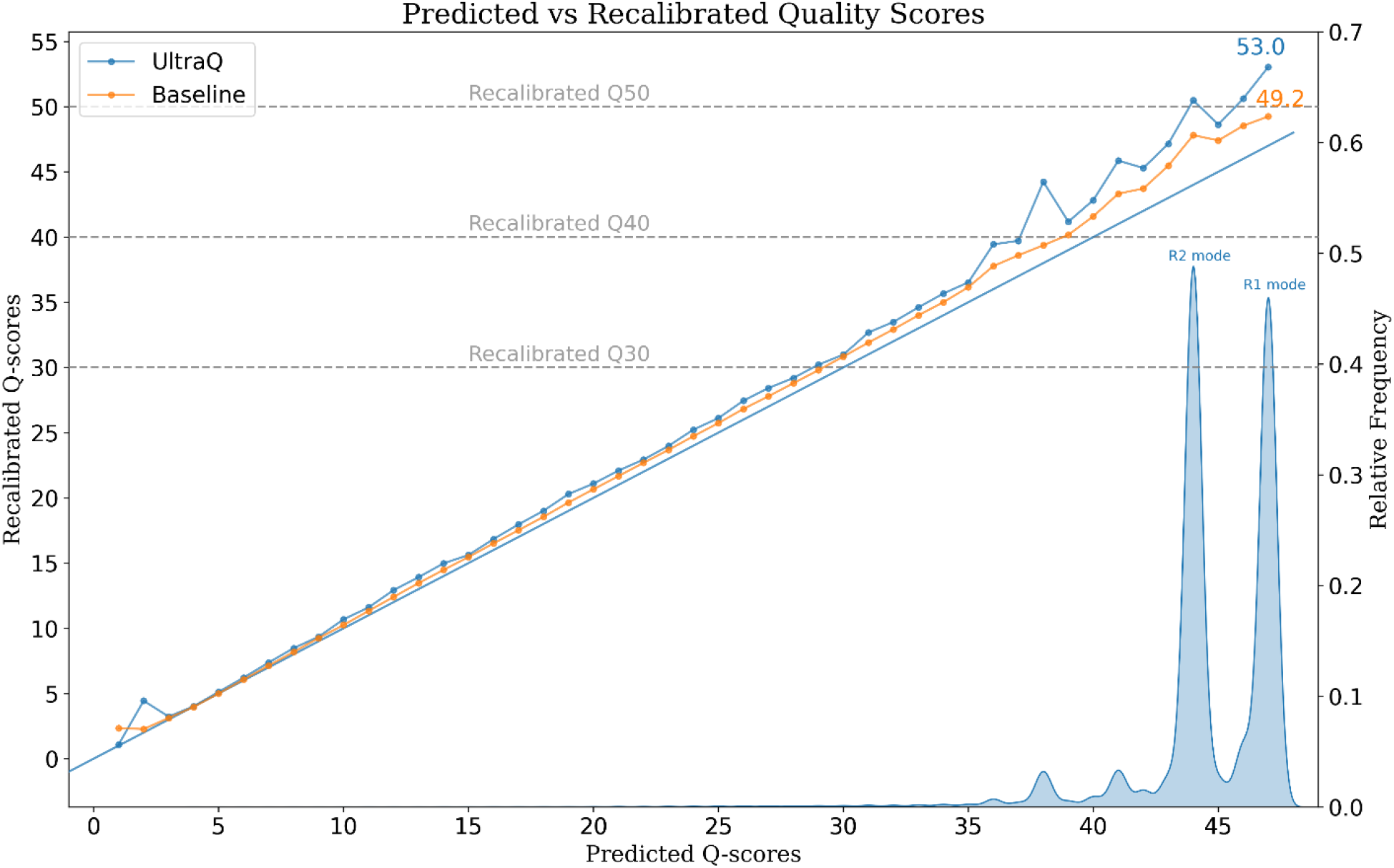
Recalibrated quality scores for a baseline and UltraQ sequencing run. Blue and orange lines show predicted vs recalibrated quality scores. The UltraQ run recalibrates above Q50 in the highest bins. The density plots below the line show the distribution of data among the predicted quality bins. The large peaks are the modes of R1 and R2.

Finally, we applied the UltraQ sequencing to a human whole genome sequencing library. Rather than looking at recalibration of quality scores, where analysis artifacts play a large role, we compare mismatch rates. Mismatch rates include not only errors but also human variation. Figure 5 shows the benefits of the UltraQ recipe translate to human data.

**Figure 5:**
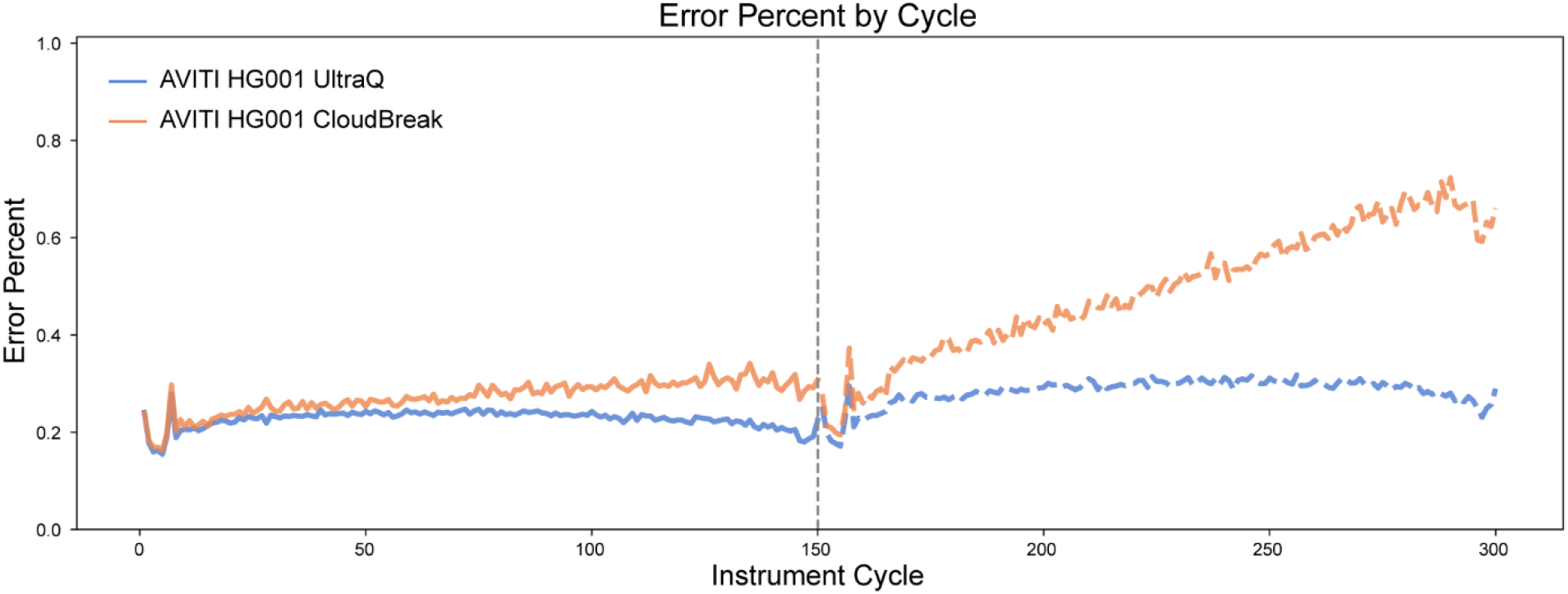
Mismatch rate comparison of baseline (Cloudbreak chemistry) and UltraQ recipe. Mismatch rate includes variation. Error components are dominated by low quality sequencing errors. However, the UltraQ recipe clearly outperforms the baseline, indicating that the benefits of the recipe generalize to human sequencing.

## Discussion

This study shows that a high fraction of Q50 data can be achieved with ABC sequencing through a combination of methods that target sequencing errors, library preparation errors, and analysis artifacts. Improvements to the sequencing recipe led to a higher fraction of data assigned to high quality scores. Conditioned on a high-quality score, library preparation errors and analysis artifacts make up most errors. It is therefore important to focus on processes upstream and downstream of sequencing to significantly improve platform accuracy.

The current study has several important limitations. First, most of the results are derived from model systems that do not have the sequence context or analysis complexities of the human genome. Our rationale was that model systems would enable us to capture the relative error contributions of library preparation and sequencing, while limiting the confounding factors associated with analysis artifacts. Also, any improvements made to sequencing and library preparation would also benefit human sequencing (see Figure 5). In prior work, we showed the performance of ABC sequencing in homopolymers and other difficult regions^7,11^. And in future work we intend to sequence the haploid CHM13 cell line and use the associated T2T reference, to capture the sequencing contexts of the human genome while minimizing analysis artifacts.

Second, the initial results were based on very short fragment lengths, which may not be representative of standard sequencing libraries. The short fragments were crucial for enabling us to characterize errors. Although there may be some differences in sequencing quality and mapping quality of short fragments, they provide a useful development tool. The results of our optimized recipe and our conclusions regarding Q50 are established for standard length libraries (see figure 4). Third, this analysis focused on errors with a high quality score. Although lower quality errors make up the vast majority of errors in the sequencing data, the knowledge that the base calls are reduced quality is used in downstream applications to limit their impact. The optimized sequencing recipe results in more than 90% of bases exceeding Q40 before recalibration, so focusing on this subset captures most of the data that is produced.

Despite the limitations, the study proved useful in identifying and addressing important sources of error. Improvements to the sequencing recipe combined with the mitigation of library preparation errors enabled us to achieve a high percentage of Q50 data in *E. coli* sequencing and to demonstrate reduced mismatch rates in human sequencing.

## Footnotes

† Dark cycling proceeds through the incorporation of unlabeled nucleotides, skipping avidite binding and imaging portions of the sequencing cycle. Dark cycling at the beginning of a read omits the end of the fragment and slightly shifts the fragment length distribution to the left.

## Notes

### Competing Interest Statement

All authors are employees of Element Biosciences and may hold stock options.

